# Modulation of Viability, Proliferation, and Stemness by Rosmarinic Acid in Medulloblastoma Cells: Involvement of HDACs and EGFR

**DOI:** 10.1101/2023.08.10.552840

**Authors:** Alice Laschuk Herlinger, Gustavo Lovatto Michaelsen, Marialva Sinigaglia, Lívia Fratini, Gabriela Nogueira Debom, Elizandra Braganhol, Caroline Brunetto de Farias, Algemir Lunardi Brunetto, André Tesainer Brunetto, Mariane da Cunha Jaeger, Rafael Roesler

## Abstract

Medulloblastoma (MB) is a heterogeneous group of malignant pediatric brain tumors, divided into molecular groups with distinct biological features and prognosis. Currently available therapy often results in poor long-term quality of life for patients, which will be afflicted by neurological, neuropsychiatric, and emotional sequelae. Identifying novel therapeutic agents capable of targeting the tumors without jeopardizing patients’ quality of life is imperative. Rosmarinic acid (RA) is a plant-derived compound whose action against a series of diseases including cancer has been investigated, with no side effects reported so far. Previous studies have not examined whether RA has effects in MB. Here, we show RA is cytotoxic against human Daoy (IC_50_= 168 μM) and D283 (IC_50_= 334 μM) MB cells. Exposure to RA for 48 h reduced histone deacetylase 1 (HDAC1) expression while increasing H3K9 hyperacetylation, reduced epidermal growth factor (EGFR) expression, and inhibited EGFR downstream targets extracellular-regulated kinase (ERK)1/2 and AKT in Daoy cells. These modifications were accompanied by increased expression of *CDKN1A*/p21, reduced expression of *SOX2*, and a decrease in proliferative rate. Treatment with RA also reduced cancer stem cell markers expression and neurosphere size. Taken together, our findings indicate that RA can reduce cell proliferation and stemness and induce cell cycle arrest in MB cells. Mechanisms mediating these effects may include targeting HDAC1, EGFR, and ERK signaling, and promoting p21 expression, possibly through an increase in H3K9ac and AKT deactivation. RA should be further investigated as a potential anticancer agent in experimental MB.

## Introduction

Medulloblastoma (MB) is the most common malignant pediatric brain tumor, contributing significantly to the high lethality of childhood intracranial tumors (Northcott et al. 2019). This tumor is a heterogeneous group with distinct molecular and clinical characteristics. Currently, available therapy combines maximum surgical tumor resection, radiotherapy and/or chemotherapy, often leading to poor long-term quality of life for patients, like cognitive, motor, neuropsychiatric and emotional deficits (Juraschka and Taylor 2019). Therefore, searching for novel therapeutic agents capable of targeting the tumor while preserving the quality of life of patients is of paramount importance, especially among pediatric patients.

Recent efforts rely on different strategies that vary from more pleiotropic to more specific signaling pathway-directed compounds. Therefore, epigenetic modulators such as histone deacetylase inhibitors (HDACi), as well as inhibitors of different receptors such as tyrosine kinase receptors (TRK) including the epidermal growth factor receptor (EGFR), or intracellular mediators such as mitogen activated protein kinase (MAPK) and MEK, alone or combined, are currently being considered as candidate therapies for MB (Freire et al. 2023; Jaeger et al. 2013; 2020; Perla et al. 2020; Thomaz et al. 2019).

Rosmarinic acid (RA) is a phenolic compound abundant in plants, especially in the *Lamiaceae* family, such as rosemary (*Rosmarinus officinalis*) and lemongrass (*Melissa officinalis*) (Zheng and Wang 2001). It has various pharmacological and biological properties, including antitumor activities (Nadeem et al. 2019). As a food supplement, RA has already been proposed to be effective against several cancer types in preclinical studies, with no adverse events being reported (Nadeem et al. 2019). Moreover, in healthy humans, a placebo-controlled randomized study has shown that a single dose up to 500 mg has not raised any safety concern regarding clinical manifestations or changes in routine blood tests (Noguchi-Shinohara et al. 2015). Similar findings were reported by a second study, in patients presenting mild dementia due to Parkinson’s disease, who received 500 mg of RA daily for 48 weeks (Noguchi-Shinohara et al. 2020). Also, the blood-brain barrier (BBB) permeability of RA has been described (Sánchez-Martínez et al. 2022), and targeted systems for delivery of RA across the BBB are currently already being studied (Kuo and Rajesh 2017).

Studies have previously demonstrated that RA can modulate HDAC expression (Ferdousi et al. 2019; Jang et al. 2018) and EGFR activity (Balogun et al. 2021; Tumur et al. 2015), which are targets for MB therapies. Although its antitumor effects against other types of central nervous system (CNS) cancer cell lines have been demonstrated (Khan et al. 2019; Ramanauskiene et al. 2016; Şengelen and Önay-Uçar 2018), previous studies have not examined its possible effects in MB models.

Here, we aimed at investigating the effects of RA in two human cell lines representatives of the sonic hedgehog (SHH) and Group 3 molecular MB subgroups (Daoy and D283, respectively), and exploring some of the underlying molecular mechanisms.

## Materials and Methods

### Cell Culture

Human MB cell lines Daoy (HTB186™) and D283 Med (HTB185™) were obtained from the American Type Culture Collection (ATCC, Rockville, USA) and tested to confirm line identity and rule out contamination. Cells were grown and maintained in low glucose Dulbecco’s modified Eagle’s medium (DMEM; Gibco, Grand Island, USA) containing 2% (*w*/*v*) L-glutamine and 10% (*v/v*) fetal bovine serum (FBS, Gibco), and antibiotics. Cells were kept at a temperature of 37°C, a minimum relative humidity of 95%, and an atmosphere of 5% CO_2_.

### Drug Treatment

RA (Sigma-Aldrich, St Louis, USA, cat. #4033) was diluted in dimethyl sulfoxide (DMSO, Sigma-Aldrich) at the concentration of 100 mM (stock solution). Further dilutions were freshly prepared prior to use by dissolving the stock solution in DMEM/10% FBS at the concentrations described below for each experiment. Vehicle (1% DMSO) was used as negative control.

### Cell Viability

Cell viability was assessed with trypan blue cell counting (Jaeger et al. 2013). Cells were seeded at 4 x 10^3^ cells/well in DMEM/10% FBS in 96-well plates and allowed to rest overnight before treatment. The medium was replaced, and RA (100-500 μM), or 1% DMSO (negative control), or fresh medium only (untreated negative control) was added to the culture (100 μL/well). Drug dose ranges were experimentally determined based on previous studies using other cell types (Jang et al. 2018).

Cell counting was performed 48 h after treatment. The medium was removed, cells were washed with phosphate buffer saline (PBS, pH 7.4) and 50 μL of 0.25% trypsin/EDTA (Invitrogen, Wathan, USA) was added to detach cells, which were immediately counted in a hemocytometer. Experiments were performed at least three times with triplicate wells for each drug concentration.

Relative cell viability, as compared to the negative control (1% DMSO) was calculated for each drug concentration. Nonlinear regression was performed to estimate the IC_50_ for RA in each cell line. The obtained value was further confirmed by an independent experiment.

### Astrocyte Viability

Primary mouse astrocyte cultures were obtained from 2 or 3 days-old C57/Bl-6 mouse pups, as described in Schildge et al. (2013), with minor modifications. Briefly, four mouse brains were isolated and sustained in CMF buffer until they were dissected in order to remove meningeal cells and fibroblasts. Once isolated, the cortex structure from the brains were mechanically dissociated and high glucose DMEM (pH 7.6), supplemented with 10% FBS, in T75 flasks, at 37°C under 5% CO_2_ atmosphere. After seven days the mixed cortical cell astrocytes were submitted to 30 min shaking at 180 rpm to remove microglia. Subsequently, cells were incubated for another 12-14 days. Afterwards, they were seeded in 48-well plates, in a concentration of 2,000 cells/well, and allowed to grow to confluence for three days before treatment. On the third days, the medium was replaced by fresh medium containing 1% DMSO, or RA at 100, 150, 200, 250, 300, 350, and 400 μM, as well as at the IC_50_ for Daoy and D283 cells (168 and 334 μM, respectively). An untreated control which received fresh medium only, was also included. Each concentration was tested in eighth replicates. After 48h treatment, cells were detached and counted as described above. This procedure was approved by the Ethics Committee on Animal use (CEUA) of Universidade Federal de Ciências da Saúde de Porto Alegre (UFCSPA), under protocol number 653/19.

### Cell Survival

Daoy and D283 cells (5 x 10^4^ cells/well in 6-well plates) were treated with RA at 168 μM and 334 μM (2 mL/well), respectively, for 48 h before being seeded into 12-well plates (100 cells/well). After incubation for 7-9 days, cells were fixed with 100% methanol and counterstained with 0.5% crystal violet. The number and size of colonies were calculated using the ImageJ software (National Institutes of Health, NIH, Bethesda, USA) and presented as a percentage of the control treatment (1% DMSO). Experiments were performed at least three times, with triplicate wells for each treatment.

### Cell Proliferation

To calculate the proliferative potential the cumulative population doubling (CPD) was calculated every three days across 12 days, as previously described (Silva et al. 2016). After 48 h exposure to RA at 168 μM (Daoy) and 334 μM (D283) as described above, cells were seeded at the density of 2 x 10^4^ cells/well in 24-well plates, in a treatment-free medium. Doubling time was calculated using the formula PD= Ln (FN) – Ln (IN)/Ln (2), where IN (initial number) is the number of cells plating in each passage and FN (final number) is the number of cells obtained in every counting. Experiments were performed at least three times, with triplicate wells for each treatment.

### Neurosphere Formation

Neurosphere formation was used as an experimental assay to evaluate cancer stem cell (CSC) proliferation (Nör et al. 2013) as previously described (Jaeger et al. 2020). Daoy and D283 cells were seeded (500 cells/well) in 24-well plates previously coated with 1% agarose. The cells were cultured in a serum-free sphere-induction medium, composed of DMEM/F12 supplemented with 20 ng/mL epidermal growth factor (Sigma-Aldrich), 20 ng/mL basic fibroblast growth factor (Sigma-Aldrich), B-27 supplement 1 × (Gibco), N-2 supplement 0.5 × (Gibco), 50 μg/mL bovine serum albumin (Sigma-Aldrich) and antibiotics. The RA treatment (500 μL of 168 μM and 334 μM for Daoy and D283, respectively) was added on the first day of sphere induction, on the third day 500 μL of the sphere-induction medium was added per well, and the induction protocol continued until the fifth day. Sphere photomicrographs were captured on the fifth day of induction under an inverted microscope (Leica Microsystems, Wetzlar, Germany) at a 10 × magnification. The area of the spheres was measured using the ImageJ software and is represented as the percentage of the control treatment (1% DMSO). Experiments were performed at least three times, with triplicate wells for each treatment.

### Quantitative Reverse Transcription Polymerase Chain Reaction (qPCR)

Total RNA from MB cells treated with RA for 48 h was extracted using the SV Total RNA Isolation System kit (Promega, Madison, USA), following the manufacturer’s instructions, and quantified in a NanoDrop (Thermo Fisher Scientific, Waltham, USA). The cDNA was obtained using GoScript Reverse System (Promega) from 200 ng RNA, according to the manufacturer’s instructions. PowerUp SYBR Green Master Mix (Thermo Fisher Scientific) was used to quantify the mRNA, using 10 ng cDNA, expression levels of target genes: *PROM1* (primer sense 5’-GGC AAA TCA CCA GGT AAG AAC-3’, primer anti-sense 5’-AAC GCC TTG TCC TTG GTA G-3’)*, CDKN1A* (primer sense 5’-ACT CTC AGG GTC GAA AAC GG-3’, primer anti-sense 5’-CTT CCT GTG GGC GGA TTA GG-3’)*, SOX2* (primer sense 5’-CAG CTC GCA GAC CTA CAT GA-3’, primer anti-sense 5’-GGG AGG AAG AGG TAA CCA CAG-3’), *NANOG* (primer sense 5’-AGC TAC AAA CAG GTG AAG ACC-3’, primer anti-sense 5’-GTG GTA GGA AGA GTA AAG GCT G-3’), *BMI1* (primer sense 5’-TGC TTT CTG GAG GGT ACT TC-3’, primer anti-sense 5’-GTC TGG TCT TGT GAA CTT GGA-3’), *POU5F1* (primer sense 5’-AGT GAG AGG CAA CCT GGA GA-3’, primer anti-sense 5’-ACA CTC GGA CCA CAT CCT TC-3’). Expression of *ACTB* (primer sense 5’-AAC TGG AAC GGT GAA GGT G-3’, primer anti-sense 5’-AGA GAA GTG GGG TGG CTT TT-3’) was measured as an internal control. Samples for triplicate experiments were analyzed, with three technical replicates per sample per target.

### Western Blot

Cells were lysed after RA treatment for 48h with 1X lysis buffer (Cell Lysis Buffer 10 ×, Cell Signaling Technology, Danvers, USA), with protease and phosphatase inhibitors. Proteins were quantified using the Bradford protein assay (Thermo Fisher Scientific). For blotting, 20 μg of protein were separated by SDS-PAGE and transferred onto a PVDF membrane. After 1h blocking (5% skim powder milk in TTBS), membranes were washed with TTBS and incubated overnight at 4 °C with primary antibodies: HDAC1 (1:1,000; Cell Signaling, #2062), HDAC2 (1:1,000; Cell Signaling, #2540), histone H3K9ac (1:500; Abcam, Cambridge, UK, ab10812), histone H3 (1:1,000; Abcam, ab1791); EGFR (1:1,000; Cell Signaling, #4267), pERK1/2 (1:1,000; Cell Signaling, #9101), ERK1/2 (1:1,000; Cell Signaling, #4695), pAKT (Thr308) (1:1,000; Cell Signaling, #13038), AKT (1:1,000; Cell Signaling, #9272), p21 (1:1,000; Santa Cruz Biotechnology, Dallas, USA, sc-397), and β-actin (1:5,000; Abcam, ab6276). Membranes were washed with TTBS and incubated for 1 h at room temperature with isotype specific secondary antibodies anti-mouse IgG (1:10,000; Sigma-Aldrich, A4416) and anti-rabbit IgG (1:80,000; Sigma-Aldrich, A0545). Detection was performed using Millipore Immobilon Western Chemiluminescent HRP Substrate (Thermo Fisher Scientific). Images were captured in an I Bright Imaging System (Invitrogen). Samples for triplicate experiments were analyzed.

### Differential Gene Expression

We analyzed mRNA expression data of three Daoy and three D283 samples available in the dataset GSE77947, obtained from the Gene Expression Omnibus public genomic data repository. Data quality control and normalization were performed using the R software environment v.3.4.4 with the packages “arrayQualityMetrics” and “Oligo”, respectively. Transcripts were considered to be differentially expressed between cell lines when FDR < 0.05 and -1.1 < logFC > 1.1.

### Survival Analysis

The MB dataset GSE85217 (Affymetrix Human Gene 1.1 ST Array, GPL22286) contains 763 gene expression samples and was acquired from Gene Expression Omnibus (GEO) (Edgar et al. 2002). Additional data as overall survival and molecular subgroup was retrieved from the corresponding publication (Cavalli et al. 2017). Only 172 samples of the SHH subgroup with overall survival data were kept for survival analysis. Two outlier cases detected with the quality control R/Bioconductor package “arrayQualityMetrics” were excluded (Kauffmann et al. 2009). Afterward, normalization and processing were conducted with the “oligo” R/Bioconductor package (Carvalho and Irizarry 2010). Patients with high or low EGFR expression were divided with the “maximally selected rank statistics method” (Lausen and Schumacher 1992), their Kaplan-Meier curves estimated and compared utilizing the log-rank test followed by Bonferroni correction.

### Statistical Analysis

Data are shown as mean ± standard deviation (SD). Statistical analysis was performed by paired or unpaired two-tailed Student’s T-test, one-way analysis of variance (ANOVA), or repeated measures ANOVA, followed by Bonferroni post hoc tests for multiple comparisons. Experiments were replicated at least three times; *p* values under 0.05 were considered to indicate statistical significance. The GraphPad Prism 8 software (GraphPad Software) was used for all analyses.

## Results

### RA is Cytotoxic Against MB Cells

RA significantly reduced the number of viable Daoy and D283 cells (**Fig. 1a**, **1b**). The cell lines showed different sensitivity to RA. For Daoy cells, the tested RA concentration ranged between 100 and 250 μM, with reduced viable cell number being observed at the doses of 175 μM (*p* < 0.01), 200 μM (*p* < 0.01), 225 μM (*p* < 0.001), and 250 μM (*p* < 0.001) (**Fig. 1a**). In contrast, reaching similar effects in D283 cells required drug concentrations between 250 and 500 μM, and a reduction in the number of viable cells was observed for the doses of 350 μM (*p* < 0.05); 400 μM, 450 μM and 500 μM (*p* < 0.001 for all concentrations) (**Fig. 1c**). In fact, the estimated IC_50_ for RA in Daoy and D283 cells was 168 μM (*p* = 0.029) and 334 μM (*p* = 0.009), respectively. These values were confirmed by additional independent experiments (**Fig. 1b**, **1d**), and the determined concentrations were used in the experiments that followed. Of note 1% DMSO *per se* did not significantly affect the viability of cells (**Fig. 1a and 1c**). Moreover, when RA was tested against primary mouse astrocyte cultures, it did not demonstrate toxicity towards these normal brain cells (**Fig. 1e and 1f**).

**Fig 1.**
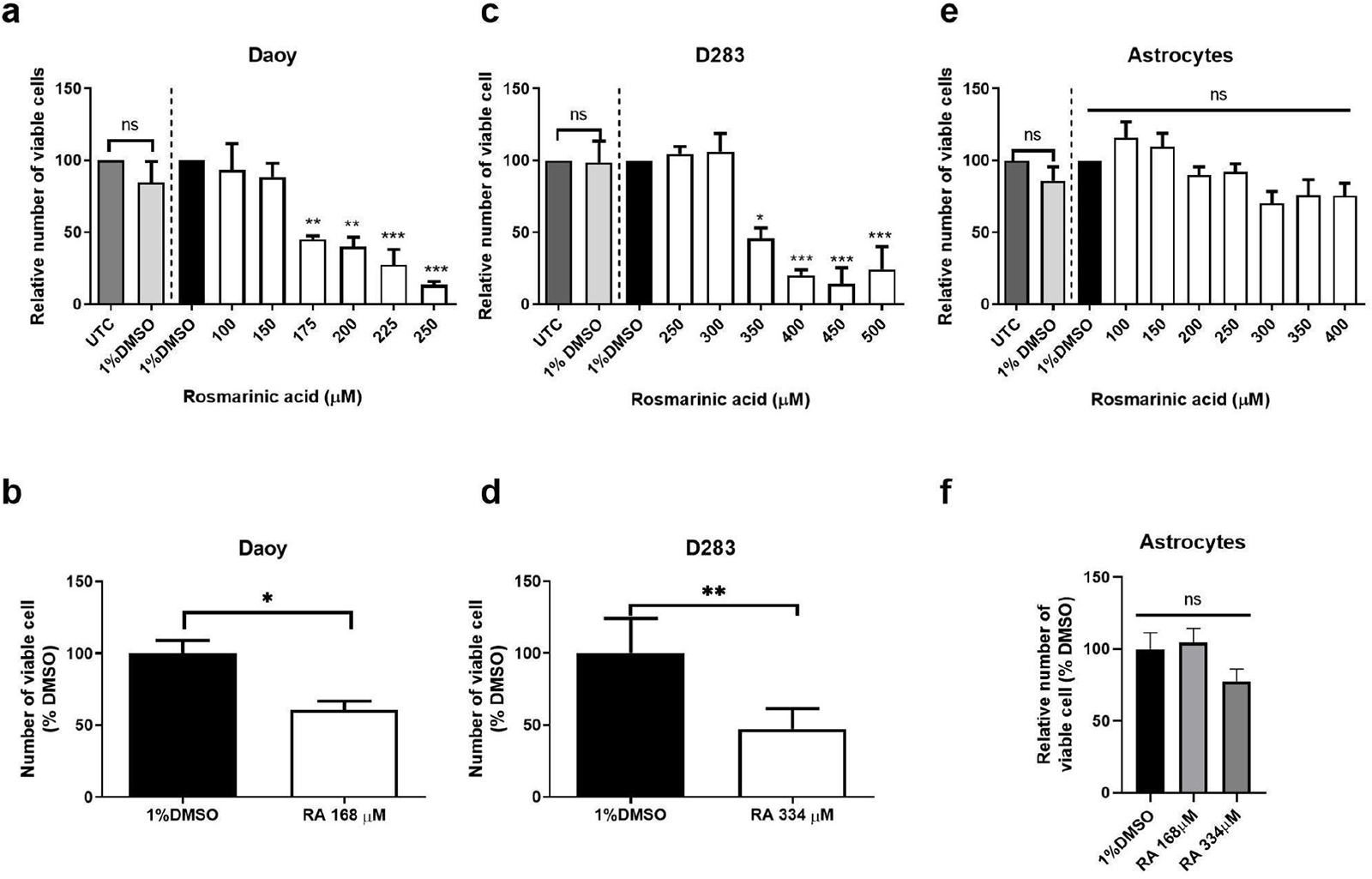
Dose dependent cytotoxic effect of RA in MB cells. A series dilution of RA was tested against (**a)** Daoy and (**c)** D283 MB cells, and primary mouse astrocytes (**e**). Cells received fresh medium (untreated control, UTC) or were treated with RA or 1% DMSO (vehicle) for 48 h prior cell viability assessment by trypan blue exclusion. Data are expressed as mean ± SD of the relative number of viable cells, as compared to controls, in three independent experiments. One-way ANOVA with Bonferroni post-hoc tests were performed to determine statistically significant differences. A nonlinear regression was performed to estimate the IC_50_ for each cell line, resulting in 168 μM for Daoy (**c**) and 334 μM for D283 cells (**d**). Primary mouse astrocytes were also treated with the estimated IC_50_ for Daoy and D283 (**f**). Data for the IC_50_ experiments are presented as mean ± SD relative number of viable cells, and the Student’s T test was performed; * *p* < 0.05; ** *p* < 0.01; and *** *p* < 0.001 compared to controls.

### RA Reduces the Area, but not the Number of Cell Colonies

We then examined the effect of RA on the ability of cells to form colonies. Therefore, after being exposed for 48 h to RA at the IC_50_ for each cell line, cells were reseeded at low density and allowed to grow for 7-9 days. As shown in **Fig. 2a** and **2b**, treatment with RA did not affect the number of Daoy colonies; however, significantly reduced the area of the formed ones (*p* < 0.001; **Fig. 2b**). In contrast, RA affected neither the number nor the area of colonies formed by D283 cells (**Fig. 2d, 2e**). These findings suggest that, in Daoy cells, RA may not prevent colony formation, but might nonetheless interfere with cell proliferation, given that the area of colonies was reduced following RA treatment.

### RA Decreases Cell Proliferation

To assess the effect of RA on the proliferation potential, we evaluated CPD. Daoy cells treated with 168 μM RA for 48 h showed a reduced CPD across 12 days (**Fig. 2c**), with an estimated population doubling time of 4.1 days (98.6 h) following RA exposure, compared to 2.5 days (59.3 h) for the control group (*p* < 0.05). D283 cells, in turn, showed a reduced CPD in response to RA only on the 8th day (*p* < 0.05; **Fig. 2f**). Although there was an apparent increase in the estimated population doubling time from 1.8 days (42.5 h) in the control group to 2.2 days (51.3 h) in RA-treated cells, this difference did not reach statistical significance (*p* = 0.12). These results confirm that RA reduces the proliferation potential of Daoy cells, but has only a limited effect in D283 cells.

**Fig 2.**
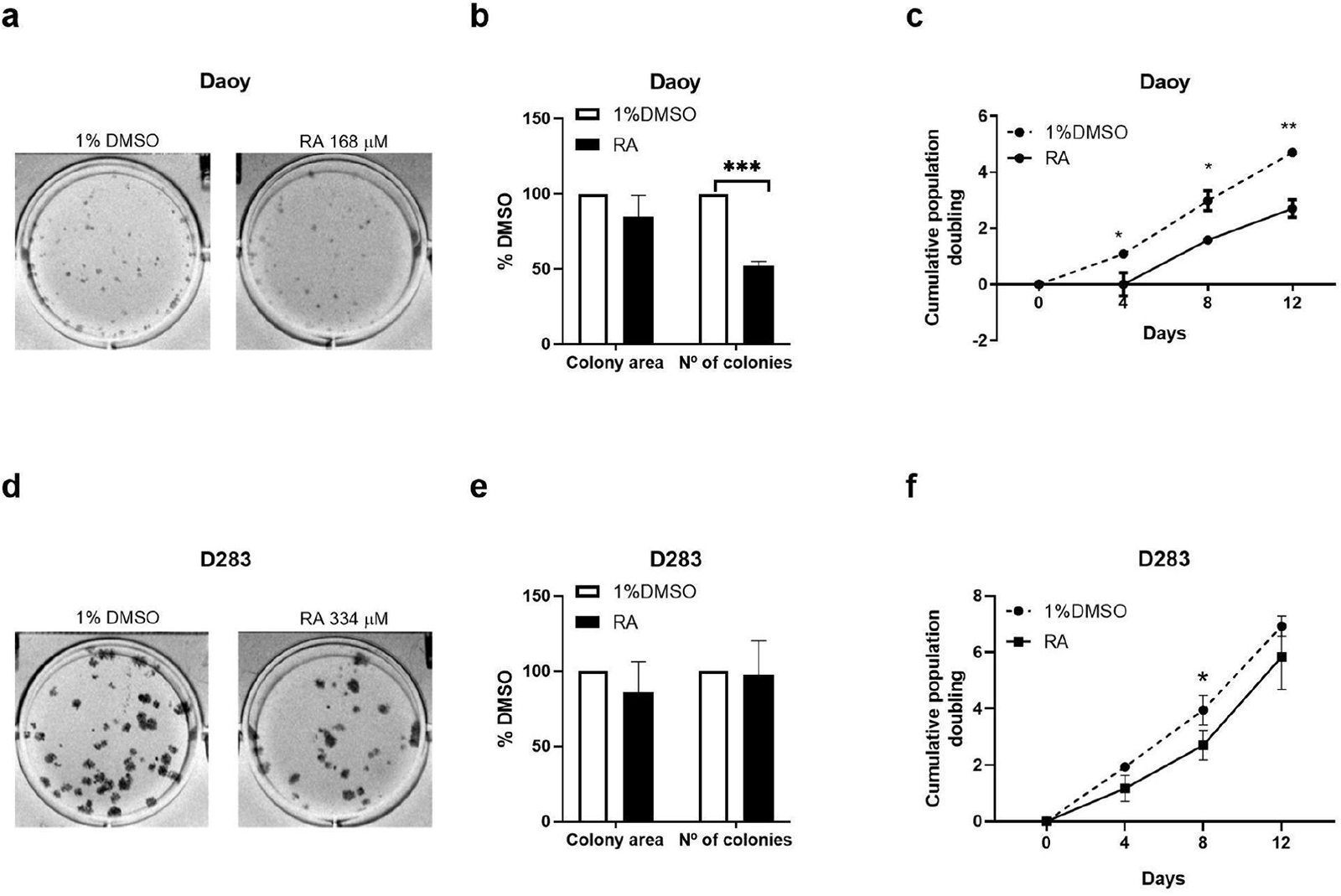
RA effects on colony formation and proliferation rate of MB cells. Daoy and D283 cells were treated with RA at 168 μM and 334 μM, respectively, or 1% DMSO (control) for 48 h, prior to clonogenicity assessment. Representative images of colonies formed from **(a)** Daoy and **(d)** D283 cells. The average number of colonies formed by Daoy and D283 cells, as well as the average colony area (**b** and **e** for Daoy and D283, respectively) were calculated relative to control cells. Data are presented as mean ± SD relative number or area of colonies, and Student’s T tests were performed for comparisons between treated and control cells; *** *p* < 0.001. For evaluation of the proliferation rate Daoy and D283 cells were treated with RA or 1% DMSO (control) for 48 h prior to proliferative rate assessment. Cells were counted and reseeded at days 4, 8 and 12. The cell counts were used to estimate the cumulative population doubling (CPD) for **(c)** Daoy and **(f)** D283 cells. Data are expressed as mean CPD ± SD. Repeated measures One-way ANOVA with Bonferroni post-tests were performed for comparisons between groups; \**p* < 0.05; and ** *p* < 0.01.

### RA Reduces the Stem Cell-Like Phenotype of MB cells

Next, we investigated the effect of RA on the stem cell-like phenotype of MB cells. Upon treatment with RA at its IC_50_, a reduction in neurosphere area was observed for both Daoy-(*p* < 0.05) and D283-derived (*p* < 0.01) spheres (**Fig. 3a-c**). In addition, the mRNA expression of the cancer stem cell (CSC) markers were reduced. Whereas *PROM1* (CD133) was reduced in both cells (*p* < 0.05 for Daoy, and *p* < 0.001 for D283), *NANOG* was downregulated exclusively in Daoy cells (*p* < 0.01), and *POU5F1*/Oct-4, in D283 cells (*p* < 0.0001). Although the reduction in the sphere area was similar between cell lines, the reduction on the expression of the CSC markers was much less profound in Daoy cells, with about 30% reduction in *PROM1* and 20% in *NANOG* expression (**Fig. 3d**), compared to a nearly 70% reduction on *PROM1* and *POU5F1* mRNA expression in D283 cells (**Fig. 3e**).

**Fig 3.**
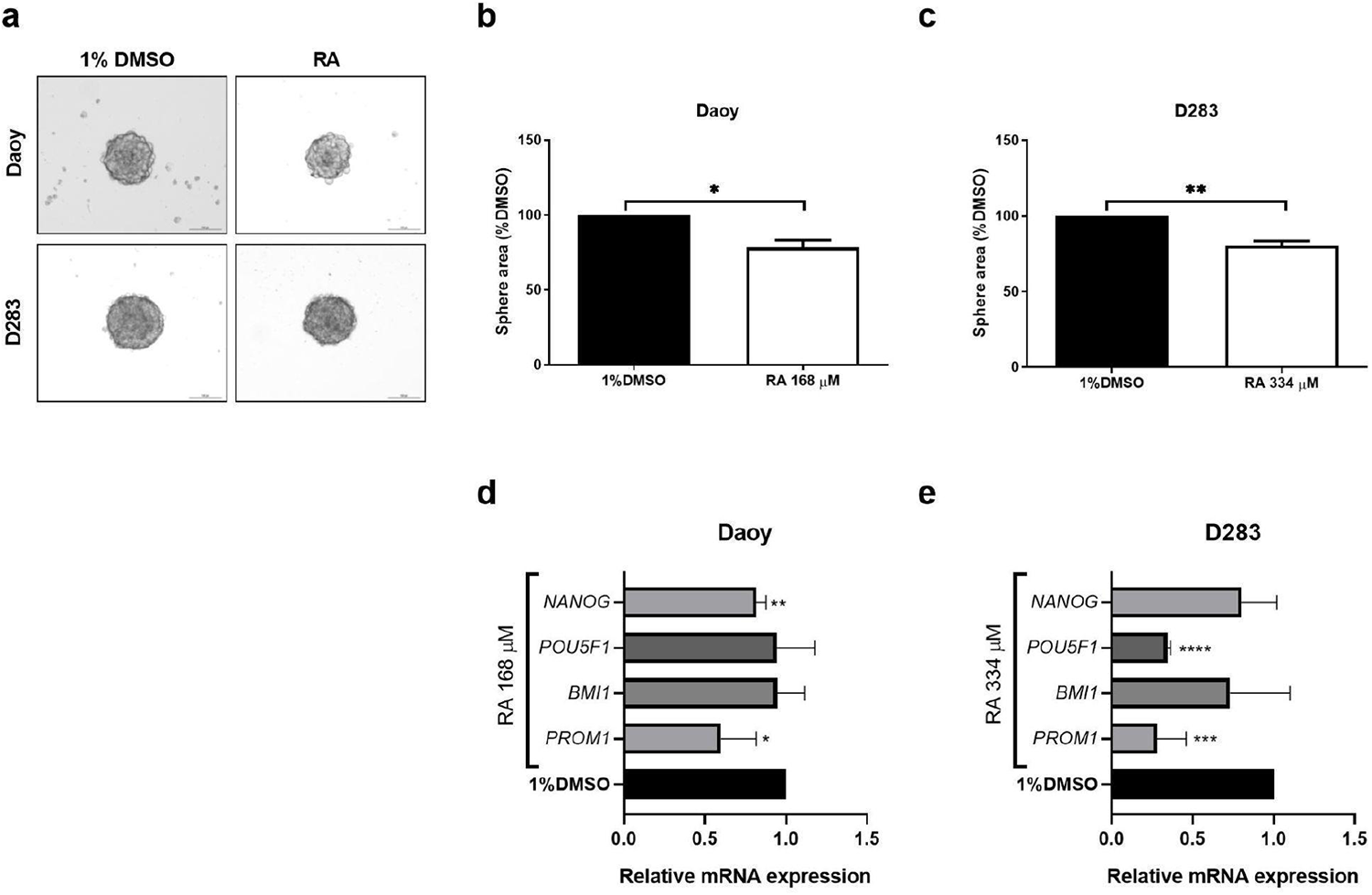
RA modulates the stem cell-like phenotype in MB cells. (**a)** Representative images of spheres formed in Daoy and D283 MB cell cultures after 5 days of induction in the presence of 1% DMSO or RA at 168 and 334 μM, respectively. Average sphere area was calculated relative to DMSO-treated **(b)** Daoy and **(c)** D283 cells. Relative mRNA levels of the stemness markers *NANOG*, *POU5F1*, *BMI1* and *PROM1* after 48 h treatment with RA of **(d)** Daoy and **(e)** D283. Data are expressed as mean ± SD of three independent experiments, and Student’s T test were performed for comparison between treated and control cells; * *p* < 0.05; ** *p* < 0.01; *** *p* < 0.001; and \*\*\*\**p* < 0.0001.

### RA Downregulates HDAC1 and Increases H3K9ac in Daoy Cells

RA has been previously shown to modulate the expression of HDACs (Ferdousi et al. 2019; Jang et al. 2018), which are increasingly being considered as therapeutic targets for MB (Perla et al. 2020). Therefore, we sought to investigate the effect of RA on the expression of HDAC1 and HDAC2, as well as the levels of the epigenetic mark H3K9ac. Results are shown in **Fig. 4**. HDAC1 was selectively reduced by RA only in Daoy cells (*p* < 0.001; **Fig. 4b**), and this effect was associated with a discrete (2-fold) increase in H3K9ac expression (**Fig. 4c**). No change was observed in D283 cells (**Fig. 4d, 4e**). RA failed to affect HDAC2 expression (data not shown).

**Fig 4.**
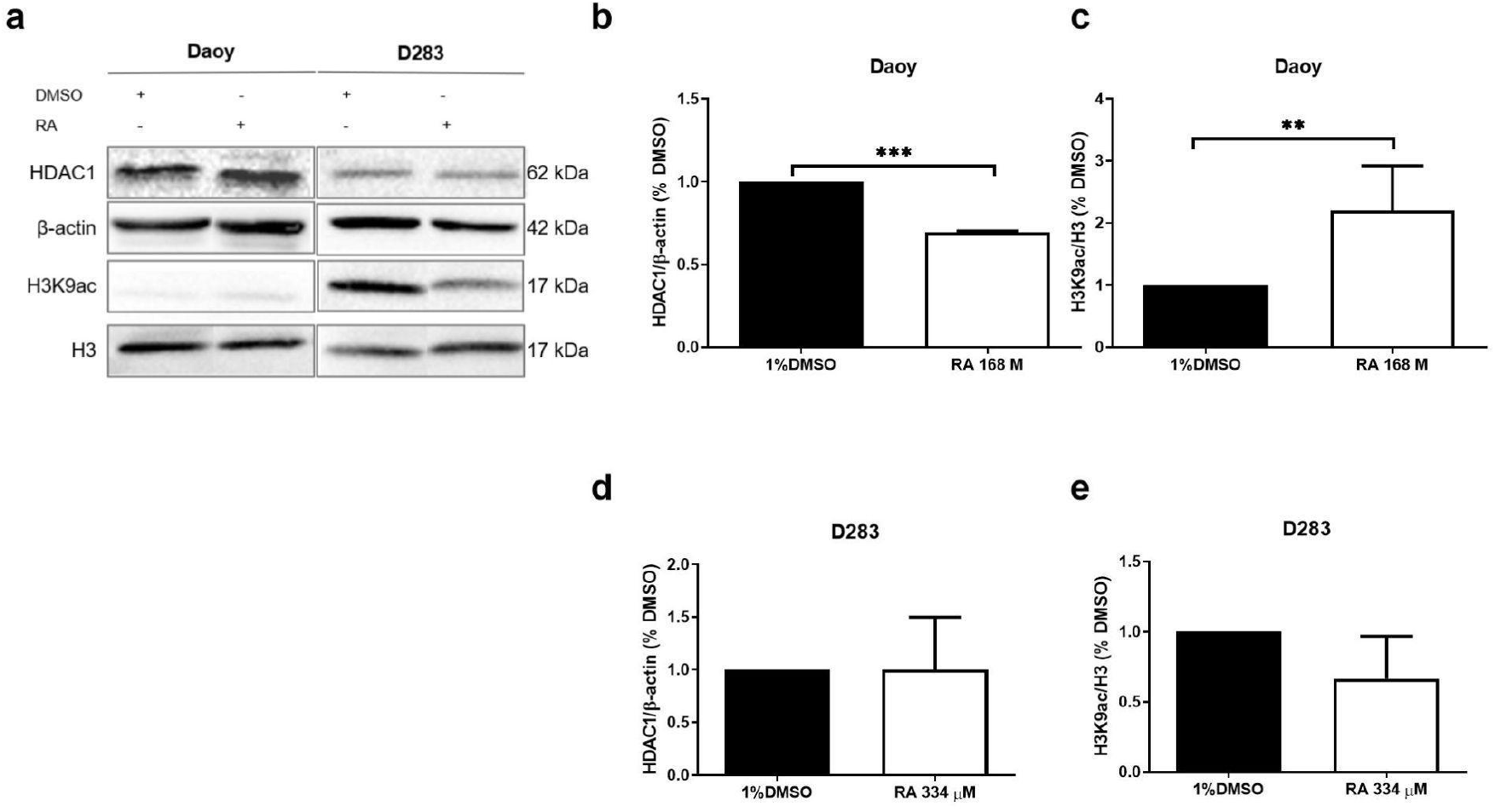
RA affects expression of HDAC1 and H3K9ac in MB cells. Daoy and D283 cells were treated with RA at 168 or 334 μM, respectively, or 1% DMSO (control) for 48 h prior to assessment of the levels of HDAC1, histone H3 acetylation at the K9 residue (H3K9ac), β-actin and histone H3 by Western blot (**a**). Expression of HDAC1 in Daoy (**b**) and D283 (**d**) cells was calculated relative to β-actin expression. Levels of H3K9ac in Daoy (**c**) and D283 (**e**) cells were calculated relative to total histone H3 expression. Data are expressed as the mean ± SD of the protein expression relative to DMSO-treated cells. Student’s T tests were performed for comparisons between treated and control cells; ** *p* < 0.01 and *** *p* < 0.001.

### RA Reduces EGFR Protein Levels in Daoy Cells

To try to further elucidate why Daoy and D283 have different responses to RA, we looked for genes differentially expressed between the cell lines treated with RA. A total of 3,287 transcripts were found to be differentially expressed between Daoy and D283 cells (data not shown). Fourteen of those genes were also modulated by RA in cancer or stem cells (**Online Resource 1**).

EGFR is a target in MB, especially in SHH group (Park et al. 2019; Jaeger et al. 2020), and was found to be overexpressed in Daoy cells when compared with D283 ones (**Fig. 5a**); despite no relation between *EGFR* expressions and patients’ overall survival was observed in a cohort of 170 SHH MB-diagnosed patients (log-rank test, *p* = 0.17; **Fig. 5b**). Nonetheless, upon exposure of Daoy cells to RA at 168 μM for 48 h, EGFR protein expression was reduced by over 50% (*p* < 0.01; **Fig. 5c and 5d**), which was not observed in D283 cells (**Fig. 5e**).

**Fig 5.**
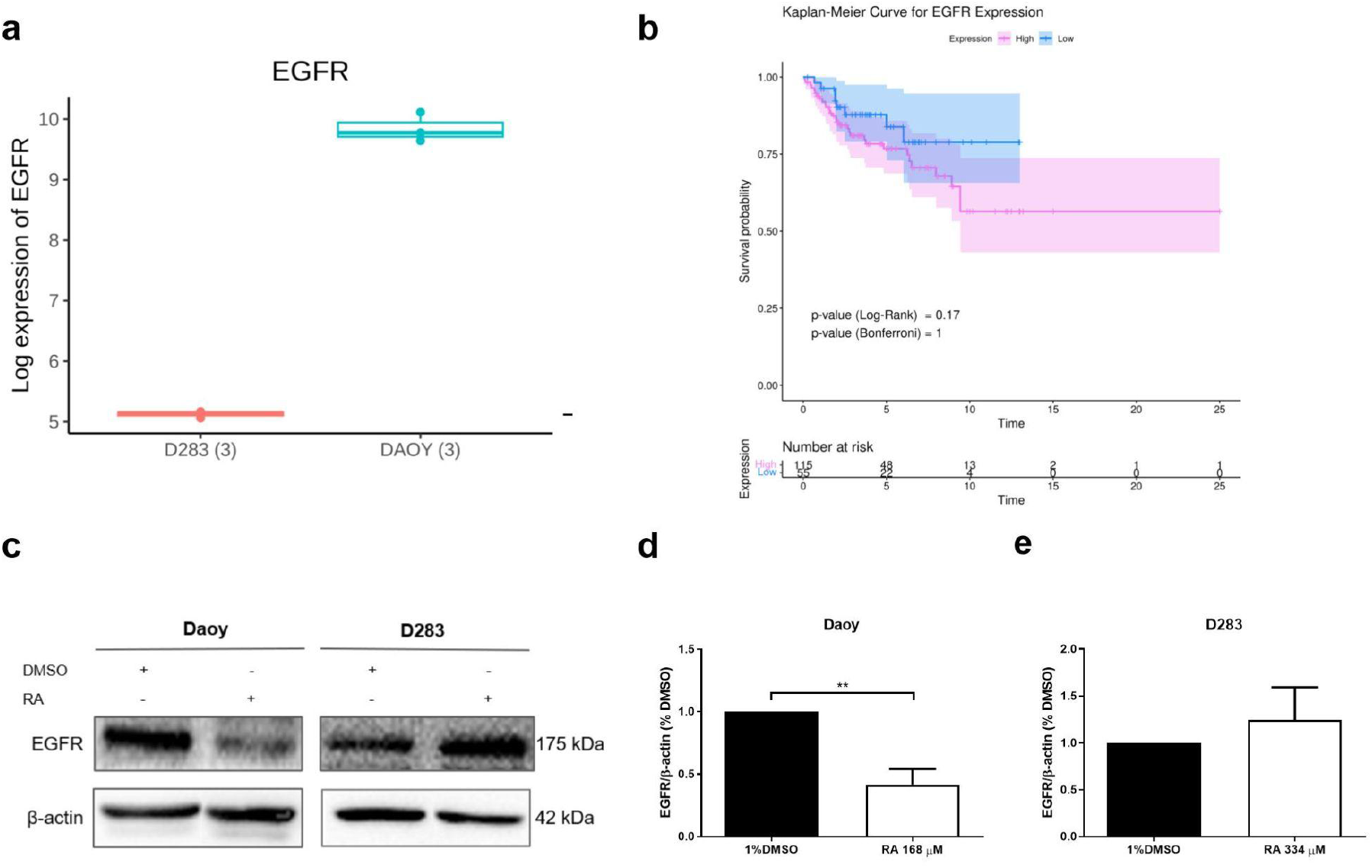
RA affects EGFR expression in MB cells. **(a)** In silico differential expression analysis. EGFR mRNA is overexpressed in Daoy cells (*n* = 3) compared to D283 cells (*n* = 3). **(b)** Overall survival and *EGFR* expression of patients with MB tumors belonging to molecular group SHH (n = 170, log-rank test, *p* = 0.17). For Western blot, Daoy and D283 cells were treated with RA at 168 or 334 μM, respectively, or 1% DMSO (control) for 48 h prior to assessment of levels of EGFR and β-actin (**c**). EGFR content in **(d)** Daoy and **(e)** D283 cells was calculated relative to β-actin expression. Data are expressed as the mean ± SD of the protein expression relative to controls. Student’s T tests were performed for comparisons between treated and control cells; ** *p* < 0.01.

### RA Disrupts EGFR Signaling in Daoy Cells

As EGFR was selectively reduced by RA in Daoy cells, we sought to investigate whether the EGFR downstream signaling pathways were also disrupted, and therefore would provide a candidate mechanism for the actions of RA. As shown in **Fig. 6**, decrease in EGFR by RA was accompanied by a reduction of nearly 50% on the levels of phosphorylated AKT at the Thr308 residue, which is phosphorylated by the upstream PDK1 (*p* < 0.01; **Fig. 6a**, **6b**), as well as phosphorylated extracellular-regulated kinase (ERK) 1/2 (*p* < 0.001; **Fig. 6c**). In agreement with these findings, *CDKN1A*/p21, a negative regulator of cell cycle progression, which has its subcellular localization controlled by AKT (Zhou et al. 2001; Li et al. 2002), was overexpressed in response to RA (*p* < 0.05; **Fig. 6d**, **6e**). Similarly, *SOX2*, also a downstream target of the EGFR/AKT and EGFR/ERK axes, was downregulated following RA treatment (*p* < 0.01; **Fig. 6f**). Expression of neither *CDK1A*/p21 nor *SOX2* was changed by RA in D283 cells (data not shown).

**Fig 6.**
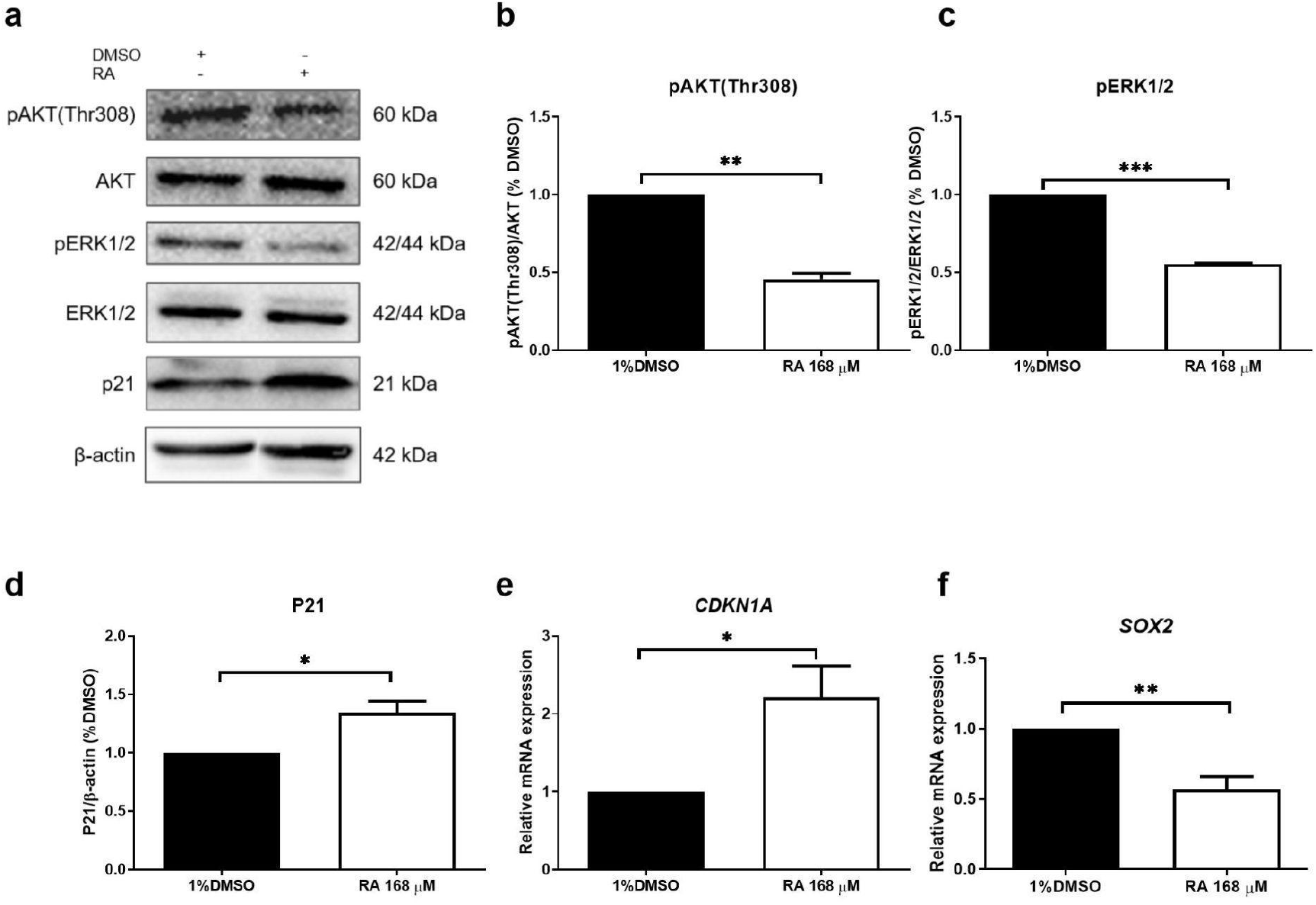
RA effects on the activation of EGFR downstream targets. Daoy cells were treated with RA at 168 μM or 1% DMSO for 48 h prior to expression levels assessment. **(a)** Protein levels of phosphorylated AKT (Thr308), total AKT, pERK1/2, total ERK1/2, p21 and β-actin were assessed by Western blot. Mean expression of pAKT(Thr308) relative to total AKT (**b**), pERK1/2 relative to total ERK1/2 (**c**), and p21 relative to β-actin (**d**) was calculated. mRNA levels of **(e)** *CDKN1A* (which encodes p21) and **(f)** *SOX2* were assessed by qPCR and calculated relative to *ACTB*. Data are expressed as mean ± SD of expression relative to control cells. Student’s T tests were performed for comparisons between treated and control cells: * *p* < 0.05; ** *p* < 0.01; and *** *p* < 0.001.

## Discussion

MBs are a group of tumors that correspond to over 60% of pediatric intracranial embryonic tumors, which are responsible for 25% of deaths related to pediatric cancer (Jones et al. 2012). Apart from the burden of the disease, the currently available therapy is drastic and associated with a poor long-term quality of life for patients reaching a cure. Neurological, neuropsychiatric and emotional disorders are common among this group of childhood cancer survivors (Juraschka and Taylor 2019), making the identification of novel therapeutic compounds capable of targeting the tumor without jeopardizing the patients’ quality of life imperative. In this scenario, RA stands out as a potential therapeutic for MB, as it has already been reported to act on several cellular targets which are relevant for MB treatment, and its use as a food supplement has suggested some efficacy without severe adverse effects in preclinical (Nadeem et al. 2019) and clinical studies (Noguchi-Shinohara et al., 2015; 2020). Here, we show for the first time the cytotoxic effect of RA against two MB cell lines, and that it is able to interfere in cellular processes crucial for MB progression. Moreover, we demonstrate that distinct molecular mechanisms seem to be involved in this antitumor effect in MB cells representative of SSH group and Group 3 tumors.

Although we observed a cytotoxic action of RA against both cell lines, Group 3 D283 cells (IC_50_=334 μM) were more resistant than SHH Daoy cells . In fact, D283 cells needed twice as much RA than Daoy cells to reach the same extent of inhibition (334 μM vs. 168 μM, respectively). Previous studies on the effect of RA in brain cancer cell lines have shown different sensibility to the compound. In the glioblastoma cell line U-87 MG, the estimated IC_50_ following 72 h RA-exposure was 40 μM (Khan et al. 2019). Using the same cell line and a 48h RA-treatment protocol, the estimated IC_50_ was 373.48 μM (Şengelen and Önay-Uçar 2018). In the rat glioblastoma cell line C6, RA IC_50_ in 24h and 48h was estimated as 290.5 and 181.3 μM, respectively (Ramanauskiene et al. 2016). Therefore, IC_50_ estimated for both Daoy and D283 are within the concentration range previously described for other CNS cancer cell lines. Of notice, when tested against primary mouse astrocytes, RA showed no toxicity, suggesting its safety towards healthy brain cells.

Despite the difference in the amount of drug required, Daoy and D283 cells showed similar responses in respect to their ability to form colonies, which was kept unchanged by the treatment, the decrease in the size of neurospheres formed, and the reduction in expression of the stemness marker *PROM1*/CD133. When we analyzed the CPD upon RA exposure, Daoy cells had a reduced CPD and longer doubling time when compared to control cells, whereas the proliferation rate of D283 cells was unaffected.

Cells giving rise to childhood cerebellar tumors closely resemble embryonic cells present during fetal development or early postnatal period (Vladoiu et al. 2019), which can be characterized by the expression of stem cell markers such as CD133 and Sox2 (Kim et al. 2003; Tamaki et al. 2002). Different from adulthood tumors, pediatric brain tumors have few somatic mutations in driver genes, but accumulate mutations in genes encoding epigenetic modulators, such as HDACs (Downing et al. 2012). These modulators have thus become promising therapeutic targets (Perla et al. 2020). RA has already been described to negatively regulate the expression of HDACs. In fact, it was shown to induce neurological differentiation of pluripotent human amnion epithelial cells through modulating a range of cellular processes, including reducing histone acetylation by downregulation of class I *HDAC1* and *HDAC3* (Ferdousi et al. 2019). Using prostate cancer cell lines, Jang et al. (2018) have shown that RA downregulates HDAC2, while promoting cell cycle arrest, reducing colony and sphere formation.

Here, we show that RA reduces HDAC1, but not HDAC2, expression in Daoy cells. HDAC1 plays an important role during early mouse development, and its siRNA-mediated knockdown leads to delayed embryonic differentiation and increased expression of p21; neither HDAC2 nor HDAC3 silencing had the same effect in mouse embryos (Ma and Shultz 2008). In fact, inhibition of HDACs strongly induces p21 transcription through several mechanisms including a reduction in HDAC1 activity (Gui et al. 2004). Another study using murine models has shown that, during embryonic neuronal differentiation, an inverse correlation between HDAC and H3K9ac expression occurs, with H3K9 hyperacetylation leading to Sox2 depletion (Vecera et al. 2017). The reduction of HDAC1 by RA in Daoy cells was accompanied by an increase in H3K9ac and p21, reduction of Sox2, and loss of proliferative potential. Thus, it is possible that, by acting as an HDAC1 inhibitor in Daoy MB cells, RA can promote p21 expression, resulting in cell cycle arrest. Through a similar mechanism, RA might lead to H3K9 hyperacetylation, Sox2 downregulation, and promote differentiation.

Aiming to identify additional mechanisms by which RA could interfere in Daoy cell proliferation without affecting D283 cells, we sought to identify differentially expressed RA target genes between the cell lines. Among the 14 genes who met these criteria, *EGFR* stood out. EGFR belongs to the ErbB family of receptor tyrosine kinases (TRKs), which, upon ligand binding, undergo homo- or heterodimerization and subsequent activation of downstream pathways, including MEK/ERK and phosphoinositide 3 kinase (PI3K)/PDK1/AKT, promoting cell proliferation and survival (Normanno et al. 2006). EGFR is the most frequently overexpressed receptor in various human tumor types (Aaronson 1991). In MB, it might serve as a marker of poor prognosis (Park et al. 2019) as well as a therapeutic target in SHH tumors (Jaeger et al. 2013). Indeed, herein we show *EGFR* expression is higher in Daoy cells as compared to D283 ones, although its expression level could not be related to SHH MB patients prognosis.

RA inhibits EGF-induced EGFR activation, leading to downregulation of PI3K/AKT and MAPK/ERK and reduction in the viability and migration of head and neck squamous cell carcinoma cell lines (Tumur et al. 2015). Also, computational and theoretical data suggest RA binds directly to EGFR, acting as a pharmacological inhibitor of the receptor (Balogun et al., 2021). We observed that RA induced a decrease of about 50% reduction in EGFR protein levels, as well as in the downstream pathways MAPK/ERK1/2 and PI3K/AKT. In neuronal precursor cells, a positive feedback loop, with reciprocal modulation of EGFR and Sox2, which depends on the EGFR-downstream pathways PI3K/AKT and MAPK/ERK 1/2, has been described (Hu et al. 2010). Therefore, we can suggest that, by directly binding to EGFR, RA blocks the positive feedback loop, resulting in the reduced expression of the receptor itself in Daoy cells, which could be responsible for the reduced cell survival.

The cyclin dependent kinase (CDK) inhibitor p21 is an important cell cycle regulator (Shamloo and Usluer 2019), which, in p53-null cancer cells, can restore the G1 checkpoint and lead to cell cycle arrest (Waye et al. 2015). To act as a cell cycle regulator, p21 must localize in the nucleus, and cytoplasmic accumulation of p21 induces cancer cell survival, proliferation and resistance to chemotherapy (Sale and Sale 2008). In glioblastoma cells, AKT directly phosphorylates p21, resulting in its accumulation in the cytoplasm and resistance to taxol (Li et al. 2002). In cells overexpressing *HER2/neu*, a member of the EGFR family, it has been shown that HER2-mediated cell growth requires AKT activation, which phosphorylates p21 and results in its accumulation in the cytoplasm (Zhou et al. 2001). In the present study, p21 was increased in Daoy cells after RA exposure, possibly due to HDAC inhibition. RA was also able to reduce EGFR and AKT. It is possible that this could prevent p21 from accumulating in the cytoplasm, inhibiting cell cycle progression.

In both cell lines, RA reduced the neurosphere area and *PROM1*/CD133. CD133 is a well-established CSC marker, involved in tumor cell proliferation, metastasis, tumorigenesis, and recurrence, as well as chemo- and radio-resistance (Jang et al. 2017). Expression of CD133 in MB cell lines has been reported by several studies, showing consistently that Daoy has low levels of CD133 expression, whereas D283 displays high expression and is proposed as an ideal CSC model (Casciati et al. 2020). Given that CD133 regulates cell death and RA can induce apoptosis in cancer cells (Huang et al. 2021; Liu et al. 2021; Messeha et al. 2020), it is possible that RA induces its cytotoxic effects at least partially by reducing CD133 in D283 cells.

## Conclusion

Our findings provide the first evidence that RA can display cytotoxic and cytostatic effects, and reduces the stemness of cultured MB cells, whereas presenting no toxic effect against healthy brain cells *in vitro*. In Daoy cells, RA affects HDAC and EGFR signaling. The ability of RA to target such pathways involved in MB progression, its demonstrated safety towards human subjects (Noguchi-Shinohara et al., 2015; 2020), as well as its ability to increase the cytotoxicity of currently used chemotherapy (Huang et al. 2021), while exerting hepatoprotection (Nadeem et al. 2019) warrants further investigations on the RA potential to MB therapy. We do acknowledge, however, the limitations of the present study, which focused on only two cell lines representative of SHH and Group 3 MB, therefore, further studies are required to better understand the mechanism of action of RA in SHH and Group 3 MB, its effects on other signaling pathways relevant to MB – such as SHH and Wingless -, its combined effect with currently used chemotherapy, and to expand the present experiments to other experimental models of MB, including animal models.

## Supporting information

Online resource

## Acknowledgements

This research was supported by the National Council for Scientific and Technological Development (CNPq, MCTI, Brazil) grants 163018/2020-0 and 101566/2022-0 to A.L.H., grants 407765/2017-4, 305647/2019-9, and 406484/2022-8 (INCT BioOncoPed) to R.R.; the Children’s Cancer Institute (ICI); and the Clinical Hospital institutional research fund (FIPE-HCPA; number 2021-0412).

## Author Contributions

A.L.H. was responsible for study conception and experimental design. A.L.H., G.L.M., M.S., L.F., G.N.B., and E.B. carried out the experiments. R.R. supervised the study. All authors contributed to statistical analysis, data interpretation, support for the study, writing and revision of this article.

## Compliance with Ethical Standards

### Conflict of Interest

The authors declare no conflicts of interest.

## Notes

### Competing Interest Statement

The authors have declared no competing interest.

### Summary of Updates

Section "Survival Analysis" and Figure 5b were revised updated to clarify that no correlation between EGFR expression and SHH MB-patients overall survival was identified.

