## Supplementary material for "Modulation of Viability, Proliferation, and Stemness by Rosmarinic Acid in Medulloblastoma Cells: Involvement of HDACs and EGFR": Online resource

**Online Resource 1** Genes modulated by RA in cancer or stem cells and overexpressed in Daoy or D283 MB cells

| Gene | logFC | *p* value | Overexpressed | Tumor/Cell Type (Reference) |
|---|---|---|---|---|
| ***Inflammation*** | | | | |
| *PLAT* | 2.75 | 3.76E-05 | Daoy | Colorectal cancer (Scheckel et al. 2008) |
| *PTGS2* | 4.25 | 1.85E-05 | Daoy | Colorectal cancer (Scheckel et al. 2008) |
| ***Adhesion, Migration, Invasion*** | | | | |
| *VCAM1* | 3.57 | 1.32E-05 | Daoy | Nasopharyngeal cancer (Lee et al. 2006) |
| *ICAM1* | 3.92 | 4.53E-06 | Daoy | Nasopharyngeal cancer (Lee et al. 2006) |
| *MMP2* | 3.09 | 1.18E-04 | Daoy | Colon carcinoma (Han et al. 2018) |
| ***Apoptosis*** | | | | |
| *TP53* | 1.57 | 3.54E-05 | Daoy | Oral carcinogenesis, glioma (Moon et al. 2010) |
| *CASP3* | 1.51 | 2.90E-03 | Daoy | Skin carcinogenesis (Sharmila and Manohanan,. 2012) |
| ***Oxidative Response*** | | | | |
| *MMP2* | 3.09 | 1.18E-04 | Daoy | Colon carcinoma (Xu et al. 2010) |
| ***Signaling Pathways*** | | | | |
| *EGFR* | 4.72 | 1.2E-08 | Daoy | Head and neck carcinoma (Tumur et al. 2015) |
| ***Proliferation*** | | | | |
| *BIRC3* | 1.91 | 3.99E-04 | Daoy | Leukemia (Moon et al. 2010) |
| *BIRC2* | 1.48 | 8.53E-04 | Daoy | Leukemia (Moon et al. 2010) |
| ***Neuronal Induction*** | | | | |
| *SOX2* | 2.35 | 4.38E-04 | Daoy | Stem cells (Ferdousi et al. 2019) |
| *FZD5* | 2.66 | 1.45E-04 | D283 | Stem cells (Ferdousi et al. 2019) |
| *RSPO1* | 2.33 | 1.21E-03 | D283 | Stem cells (Ferdousi et al. 2019) |
